# Defining the Limits of hPSC Derived Models: hPSC-MSNs Fail to Recapitulate Authentic Striatal Identity

**DOI:** 10.1101/2024.06.25.600518

**Authors:** O. J. M. Bartley, N. N. Vinh, M.J. Lelos, N. M. Williams, S. V. Precious, A. E. Rosser

## Abstract

Human pluripotent stem cells (hPSCs) are increasingly used to model human disease and as donor cells for regenerative medicine. However, the fidelity of hPSC-derived cell types remains a major concern, particularly when these cells are intended to replicate complex or region-specific subtypes, such as those required to explore and treat neurological diseases. Medium spiny neurons (MSNs), the principal projection neurons of the striatum, are one such target cell type relevant to disorders such as Huntington’s disease. While protocols for generating hPSC-derived MSNs (hPSC-MSNs) exist, the extent to which these cells faithfully recapitulate their genuine counterparts is unclear.

Here, we generated isogenic human induced pluripotent stem cells (hiPSCs) from striatal (LGE) and non-neural (fibroblast) fetal tissues, and differentiated them into MSN-like cells alongside a naïve human embryonic stem cell (hESC) line. Using DNA methylation profiling and single-cell RNA sequencing, we systematically compared the epigenetic and transcriptional features of these hPSC-MSNs to authentic fetal MSNs.

Our findings reveal persistent epigenetic signatures inherited from the tissue of origin, which influence differentiation outcomes. While LGE-derived hiPSCs retained elements of a striatal-biased methylome and yielded MSN-like cells with enhanced similarity to authentic MSNs, all hPSC-MSNs remained epigenetically and transcriptionally distinct from genuine MSNs and we identified clusters of hPSC-derived cells with aberrant or incomplete phenotypes.

These results demonstrate that even isogenic hiPSC lines exhibit variable differentiation potential due to residual epigenetic memory and protocol compatibility. We highlight the need for refined protocols and rigorous benchmarking of hPSC-derived models, particularly for regionally specified neuronal subtypes. Our study underscores the complex relationship between epigenetic status, cell lineage, protocol adaptation, and differentiation outcome.

**Paper Summary:** Human pluripotent stem cells (hPSCs) are widely used to study otherwise inaccessible human cell phenotypes. However, ensuring the molecular authenticity of hPSC-derived cell types remains critical, as differences between hPSC-derived cells and their native counterparts may impact the validity of these models. Here, medium spiny neurons (MSNs; relevant for studying basal ganglia function and disorders such as Huntington’s disease), serve as a valuable prototype for evaluating the fidelity of hPSC-derived cell types.

This study generated human induced pluripotent stem cells (hiPSCs) from developing fetal striatal tissues and fibroblasts, differentiating them into MSN-like cells alongside a human embryonic stem cell (hESC) line. Using single-cell RNA sequencing and DNA methylation analysis, we compared these hPSC-derived MSNs to authentic fetal MSNs.

Our findings reveal a significant epigenetic gap between hPSC-derived and authentic MSNs, suggesting that hPSC-MSNs do not acquire a complete and normal striatal epigenome. Additionally, while genetic expression of hPSC-MSNs was striatal-like, it was not equivalent, indicating abberant cells and a failure to reproduce an authentic phoenotype. This study provides insights into the challenges of achieving molecular authenticity in hPSC-derived cells and underscores the need for rigorous evaluation to enhance their utility in research and medicine.

## Introduction

Human pluripotent stem cells (hPSCs) offer transformative scientific potential through their capacity for modeling human development, studying disease mechanisms, and developing therapeutics. Their capacity for versatile differentiation makes them particularly valuable for studying human cells that are otherwise difficult to obtain. A key element of using hPSC derived models is assessing their successful differentiation, as any molecular or functional differences from genuine counterparts reduces the validity and worth of these cell models. However, hPSC derived cells are commonly only assayed for successful differentiation using a limited array of biomarkers, and are rarely directly compared to authentic cells. Addressing this gap is essential for accurately interpreting experimental results and for harnessing the full potential of hPSC models in both research and clinical applications.

Here, we examine medium spiny neurons (MSNs), a critical neuronal subtype in the basal ganglia circuitry, which is implicated in disorders such as Huntington’s disease (Bates et al., 2014). The successful differentiation of hPSC into MSN like phenotypes (hPSC-MSNs) has been reported by several groups (for review see Jareno et al., 2022). But there is mounting evidence that these cells are not equivalent to genuine MSNs, as they do not elicit equivalent functional recovery in transplant models (Bartley et al., 2021; Jareno et al., 2022), nor express equivalent genomic transcription patterns (Conforti et al., 2022).

Currently, it remains unclear why hPSC-MSNs are not yet equivalent to authentic phenotypes. Gene expression (and therefore cellular fate) are mediated and maintained by epigenetic mechanisms (Greenberg, & Bourc’his, 2019; Smith & Meissner, 2013; Moore, Le, & Fan, 2013), so it follows that these differences may be at least in part due to epigenetic disparities between authentic and “man-made” MSNs. Relevantly, there is evidence that early passage human induced pluripotent stem cells (hiPSC) retain elements of their previous epigenome, known as an “epigenetic memory” of their tissue of origin (Kim et al., 2010), which influences gene expression by partially replicating expression patterns of the original tissue (Table S1). Evidently, early passage hiPSCs derived from target phenotypes may be more capable of eliciting authenticity, especially compared to hiPSCs derived from off-target phenotypes, further suggesting an epigenetic component to this issue.

Here, we sought to (i) establish the current level of “authenticity” achievable in hPSC derived models of striatal MSNs, (ii) determine if this is at least in part due to epigenetic differences between hPSC-MSNs and authentic phenotypes, and (iii) to explore the possibility that epigenetically primed hPSCs may yield hPSC-MSNs with an improved phenotypic fidelity. To achieve this, we generated and characterised hPSC-MSNs from different cellular origins (“naïve” hESCs, epigenetically “primed” and epigenetically “hindered” hiPSCs) and compared these to authentic human fetal derived striatal MSNs.

## Results

### Generation and validation of hPSC-MSNs from isogenic LGE and fibroblast derived hiPSCs and hESCs

To ensure that genomic variability between hiPSC lines did not impact on our study, we generated isogenic hiPSCs from a single fetal donor using a non-integrating sendai virus to deliver reprogramming factors. We used this method to generate hiPSCs from discrete regions of the developing human fetus. First, we chose to generate hiPSCs from the lateral ganglionic eminence (LGE), the telencephalic structure that gives rise to striatal MSNs and thus most likely to produce hiPSCs epigenetically primed for differentiation into MSNs. In contrast, although dermal fibroblasts are the most commonly used cell source for hiPSC generation, they are non-neural, thus we generated an isogenic control hiPSC line from this tissue as these hiPSCs would not be epigenetically primed for a neural/striatal fate (Figure 1A).

**Figure 1.**
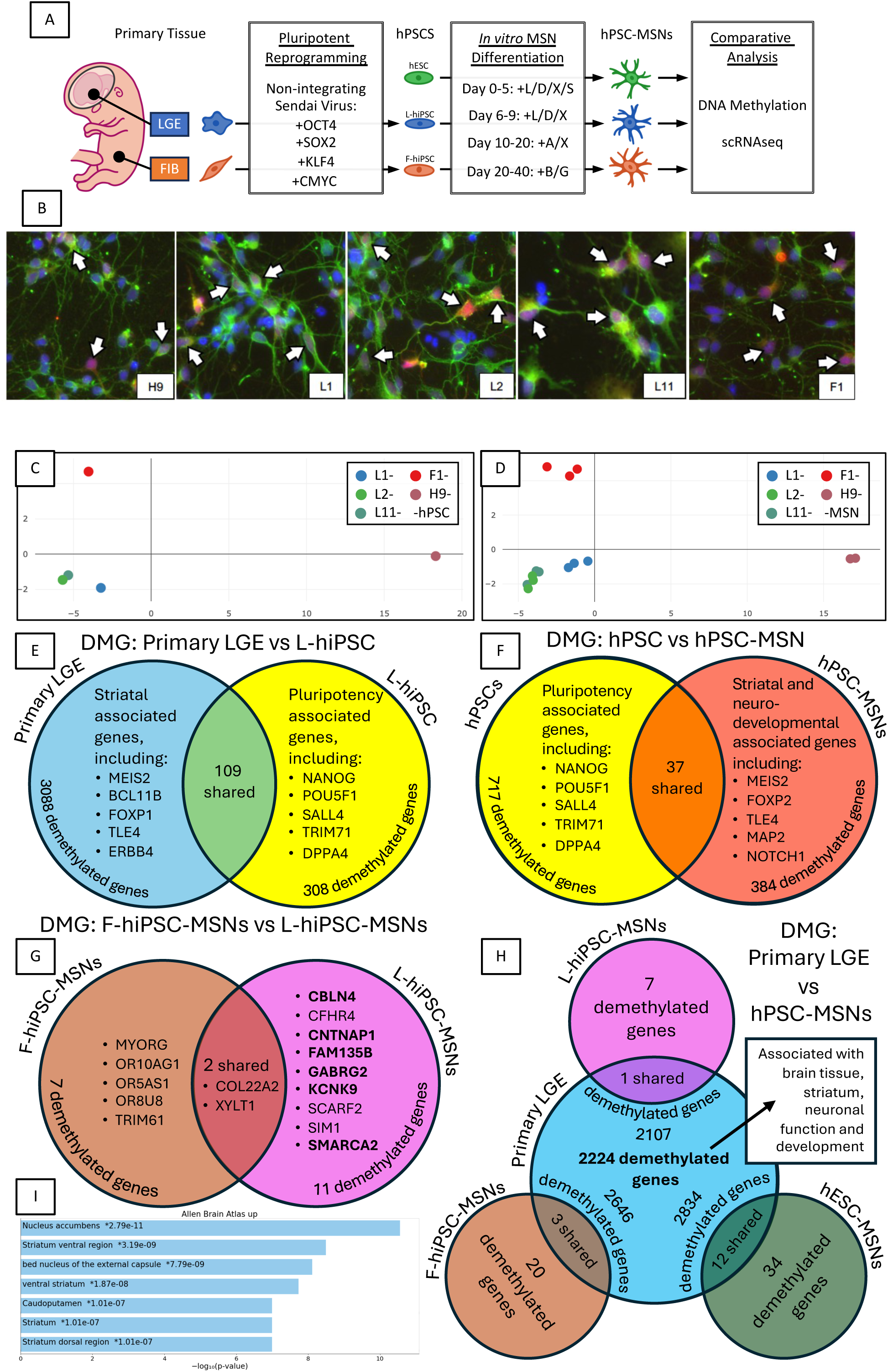
(overleaf) – Despite expressing hallmark genes, hPSC-MSNs exhibit an incomplete striatal methylome **A.** Study overview. FIB = fibroblasts; L = LDN193189; D = dorsomorphin; X = XAV939; S = SB431542; A = ActivinA; B = BDNF; G = GDNF; ICC = immunocytochemistry. **B.** Immunocytochemical staining in H9, L1, L2, L11, and F1 derived hPSC-MSNs at DIV30 for MAP2 (green) and DARPP32 (red) with Hoechst (blue). **C-D.** MDS plots of (C) pluripotent hPSCs and (D) hPSC-MSNs, produced using top 1000 most variable CPGs. **E-H.** Venn diagrams summarising significant differentially methylated genes (DMG), as demethylated genes. DMG comparisons are between (E) primary LGE (left, blue) and pluripotent L-hiPSCs (right, yellow); (F) pluripotent hPSCs (left, yellow) and differentiated hPSC-MSNs (right, orange); (G) F-hiPSC-MSNs (left, tan) and L-hiPSC-MSNs (right, pink); (H) Primary LGE (centre, blue), L-hiPSC-MSNs (top centre, pink), F-hiPSC-MSNs (bottom left, tan), and hESC-MSNs (bottom right, green). **I.** Bar chart of the most significantly enriched terms from the Allen brain atlas up database for the 2224 genes that were significantly demethylated in Primary LGE compared to hPSC-MSNs shown in H.

Colonies of hiPSCs were assessed for successful reprogramming, following which four clones were selected for further analysis; three derived from the LGE tissues (L1, L2, L11), and one from dermal fibroblasts (F1). These clones exhibited typical pluripotent cell and colony morphology, expressed critical pluripotent genes OCT4, SOX2, TRA-1-60, NANOG, and SSEA4, and were capable of producing progeny of all three germ layers (Figure S1), thus, demonstrating the successful reprogramming of these fetal tissues into hiPSCs.

We next differentiated these hiPSCs, and a separate hESC line (H9/WA09, Wicell), into hPSC-MSNs using a standardised MSN differentiation protocol (Figure 1A, MSN differentiation; adapted from Arber et al. 2015 and the RepairHD consortium). All hPSCs consistently produced neuron-rich populations (as identified by βIII-tubulin/MAP2 expression, Figure 1B, Figure S2A, S3A), which coexpressed classic MSN markers CTIP2 and DARPP32 (Figure 1B, S2B-C, S3A), indicating the successful derivation of an MSN-like phenotype across these cell lines.

### A stable tissue-specific methylation signature persists through reprogramming to pluripotency and through subsequent MSN differentiation

Having generated our hPSC lines and demonstrated their capacity to differentiate into hPSC-MSNs, we next examined the epigenome of these cells, using DNA methylation as our primary readout. Pluipotent hPSCs were sampled before (hPSC) and after differentiation (hPSC-MSN, Figure S3A) and compared to their tissue of origin and additional LGE tissues (Table S2). First, we explored whether our hPSC lines had unique methylome signatures and if this difference was maintained following differentiation into hPSC-MSNs. In their pluripotent form, hiPSCs were more similar to other hiPSCs than to hESCs, reflecting their common genomic heritage and form of pluripotency (i.e. induced; Figure S3B-C). However, we also observed that hiPSCs derived from LGE tissues were more similar to each other than to those derived from the isogenic fibroblast tissues, indicating a common residual methylation signature likely resulting from tissue of origin (Figure 1C). Importantly, this methylation pattern also persisted in the hPSC methylomes following differentiation to MSN-like cells (Figure 1D), indicating that this methylation signature is stable within these lines and has become a line-specific signature.

Analysis of differentially methylated probes (DMP) between L-and F-hiPSC-MSNs allowed us to better define this signature and explore how this could influence gene expression in MSN populations (Figure S3D). Few DMPs were observed between the LGE-derived hiPSC-MSN lines, indicating that a shared genome and tissue was the primary driver of line-specific signatures (Figure S3E). Subsequently, for further comparisons, LGE-derived hiPSC-MSNs were pooled into one group (L-hiPSC-MSN). Between L-hiPSC-MSNs and F-hiPSC-MSNs there were 437 significant DMPs, which were associated with 16 genes (Figure 1G; Table S5). Methylation of DNA commonly indicates a change in gene expression, with reduced methylation often being associated with an increase in transcription and vice versa (Bird, 2002). Of the 11 genes which exhibited reduced methylation in L-hiPSC-MSNs compared to F1-hiPSC-MSNs, five were associated with brain tissue; CNTNAP1, FAM135B, KCNK9, GABRG2 and CBLN4, and two of these (GABRG2 and CBLN4) are specifically associated with GABAergic neurons. Conversely, of the 7 genes which exhibited increased methylation in L-hiPSC-MSNs compared to F-hiPSC-MSNs, four (OR5AS1, OR8U8, OR10AG1, and MYORG) were associated with olfactory pathways and function.

Furthermore, three of the genes observed to be more methylated in F-hiPSC-MSNs compared to L-hiPSC-MSNs are down regulated in the olfactory bulb (CNTNAP1, FAM135B and SMARCA2). Notably the dorsal LGE gives rise to non-striatal cells which contribute to the olfactory bulb (Stenman et al., 2003), the expression of which are likely to be undesirable in hPSC populations desined for an MSN fate.

Overall, while the line specific differences between our hiPSC lines only affected a small number of genes, these differences do appear to be relevant to the MSN phenotype and this suggests primed LGE derived hiPSCs may be more suited to recreating the MSN epigenome compared to non-primed hiPSCs.

### Despite acquiring a striatal-like methylome, hPSC-MSNs do not recapitulate the complete methylome of authentic striatal tissues

Next, we tracked how the methylome shifted during induction to pluripotency and through differentiation into hPSC-MSNs.

Comparisons of pluripotent L-hiPSCs to LGE tissues revealed that following induction to pluripotency, L-hiPSCs were epigenetically distinct from their tissues of origin and instead clustered closer to other pluripotent cells (Figure S3B). This difference was largely driven by an increase in global genomic methylation following reprogramming, but some regions that were previously hypermethylated in original fetal tissues became hypomethylated during reprogramming (Figure S3C). We identified 60197 significantly DMPs between pluripotent L-hiPSCs and primary LGE tissues, which were associated with 3287 genes. Of these genes, 3088 became more methylated following induction to pluripotency and 308 became less methylated (109 genes exhibited both an increase and decrease in methylation across associated CpG sites, Figure 1E, Table S3). Gene enrichment analysis revealed many of the genes which accrued methylation (associated with reduced expression) following reprogramming were associated with the striatum, neuronal processes and development, whereas genes which became demethylated (associated with increased expression) were associated with pluripotent functions (Figure 1E), demonstrating a removal of striatal-specific methylome and gene expression, and restoration of a more naïve pluripotent state during induction to pluripotency.

Comparisons of hPSCs to hPSC-MSNs allowed us to characterise methylation changes following *in vitro* differentiation. Between these two cell types we identified 81627 DMPs, which were associated with 1064 genes. Of these, 717 accrued methylation during differentiation and 384 became less methylated (37 genes exhibited both an increase and decrease in methylation across associated CpG sites, Figure 1F, Table S4). As above, genes that were demethylated in pluripotent cells were strongly associated with pluripotency phenotypes, cycles and functions. Whereas, genes which were more demethylated in hPSC-MSNs were associated with striatal tissues, and biological processes related to neural development and function, including canonical MSN genes such as MEIS2, FOXP2, and TLE4 (Figure 1F). Thus indicating that our MSN differentiation protocol is specifically inducing epigenetic changes at relevant genomic regions for the MSN phenotype.

Next, to confirm whether the epigenetic signature of our LGE derived hiPSCs allowed them to produce MSNs more similar to authentic MSNs, we compared the methylome of our various hPSC-MSNs to authentic LGE tissues. Despite their varied epigenetic background, each line of hPSC-MSNs exhibited over 100,000 DMPs when compared to primary LGE, indicating large discrepancies between the methylomes of these cell populations (Tables S6-9). Examination of the associated genes revealed that 2231 genes were differentially methylated between primary LGE and hPSC-MSNs, with the vast majority (2224) exhibiting reduced methylation in primary LGE tissues compared to hPSC-MSNs (Figure 1H). These genes were strongly associated with brain tissues and striatal substructures (Figure 1I), many of which were also associated with neuronal development and activity (Figure S4).

Notably, LGE tissues comprise a heterogeneous population of cell types spanning different phenotypes and maturities, and therefore these differences may be partly driven by the variability of cell types in our primary LGE tissue group that were not present in the more focally fate-restricted hPSC-MSNs. In line with this, some enriched terms were associated with cell types that would not be expected in hPSC-MSN cultures (e.g. astrocytes, Figure S4). However, considering (i) that the majority of genes demethylated in the LGE are associated with neural and striatal phenotypes and (ii) that we identified very few (7-34) genes uniquely demethylated in hPSC-MSNs, it is reasonable to assume that this epigenetic disparity is largley driven by an incomplete acquisition of the authentic MSN methylome in hPSC-MSNs.

Overall, our data suggests that the global increase in methylation observed following induction to pluripotency has not been fully reversed following directed differentiation to an MSN-like phenotype, and that while hPSC-MSNs do exhibit some methylation changes across striatal genes in line with a “normal” striatal methylome, this process is not complete and hPSC-MSNs do not yet acquire a complete striatal methylome.

### Single cell RNA sequencing reveals hPSC-MSNs exhibit similar populations to authentic LGE-derived MSNs spanning full MSN development

To understand the biological impact of the epigenetic disparity observed above, we performed scRNAseq on four samples: (i) primary human LGE (11 pcw, genomically distinct from the hiPSCs); (ii) L2 hiPSC-MSNs; (iii) F1 hiPSC-MSNs; and (iv) H9 hESC-MSNs. Within the primary LGE we identified transcriptional phenotypes spanning normal MSN development and we sub-clustered cells of the MSN lineage for characterisation, using hallmark genes to identify specific stages of MSN development and subtypes (Figure 2 A-D, Figure S5). Specifically, we observed a cluster of cells positive for genes including HES1, SOX2, and NES, typical of apical progenitors of the LGE ventricular zone (Figure 2 D). The neighbouring cluster was positive for genes such as TOX3, ASCL1, and DLX2 which are each associated with basal progenitors of the subventricular zone of the LGE (Figure 2 D). The remaining clusters were all strongly positive for global MSN genes including MEIS2 and BCL11B, and were GABAergic as demonstrated by positive expression of GAD1 and GAD2 (Figure 2 C-D). The closest cluster to the basal progenitors was uniquely positive for POU3F1, which has previously been identified as a stage specific marker for pre-MSNs undergoing subtype fate decisions (Bocchi et al., 2021). However, POU3F1 was only expressed by a small proportion of this cluster; instead other genes such as LMO3 better defined this cluster in our analysis (Figure 2 C-D). Further assessment of this population through additional sub-clustering revealed a division into three populations which could be defined by early subtype MSN markers into (i) uncommitted progenitors, (ii) D1/EBF1+ progenitors, and (iii) D2/SIX3+ progenitors (Figure S5). Additionally, pseudotime analysis (Figure 2 B) revealed a fate decision occurring within this population which later leads to MSN subtype specification, further supporting the identification of this cluster as pre-MSN progenitors. Two further clusters expressed D1 subtype markers including ISL1 and EBF1 (Figure 2 C-D), suggesting they were of the D1 MSN lineage. Across these cells we observed a gradient increase in genes associated with mature D1-MSNs such as GRIK2, TAC1 and IKZF1, indicating these as immature and mature D1 MSNs (Figure 2 D). Finally, we observed three additional clusters which branched off from the D2/SIX3+ progenitors of the pre-MSNs. These clusters contained cells positive for early D2 progenitors such as SIX3 and SP9 (Figure 2 C), but could be differentiated by several genes, including CLMP, CXCL12, and ADARB2 (Figure 2 D). The ADARB2+ cluster also expressed TSHZ1 and ERBB4, suggestive of a D1/D2 MSN phenotype (also called eccentric MSNs, Lester, at al., 1993). Finally, we observed D1, D2, and D1/D2 MSNs could be identified by their unique expression of FOXP1 and FOXP2, D1 MSNs co-expressing both genes, D2 MSNs expressing just FOXP1, and D1/D2 MSNs expressing just FOXP2 (Figure 2 C-D).

**Figure 2.**
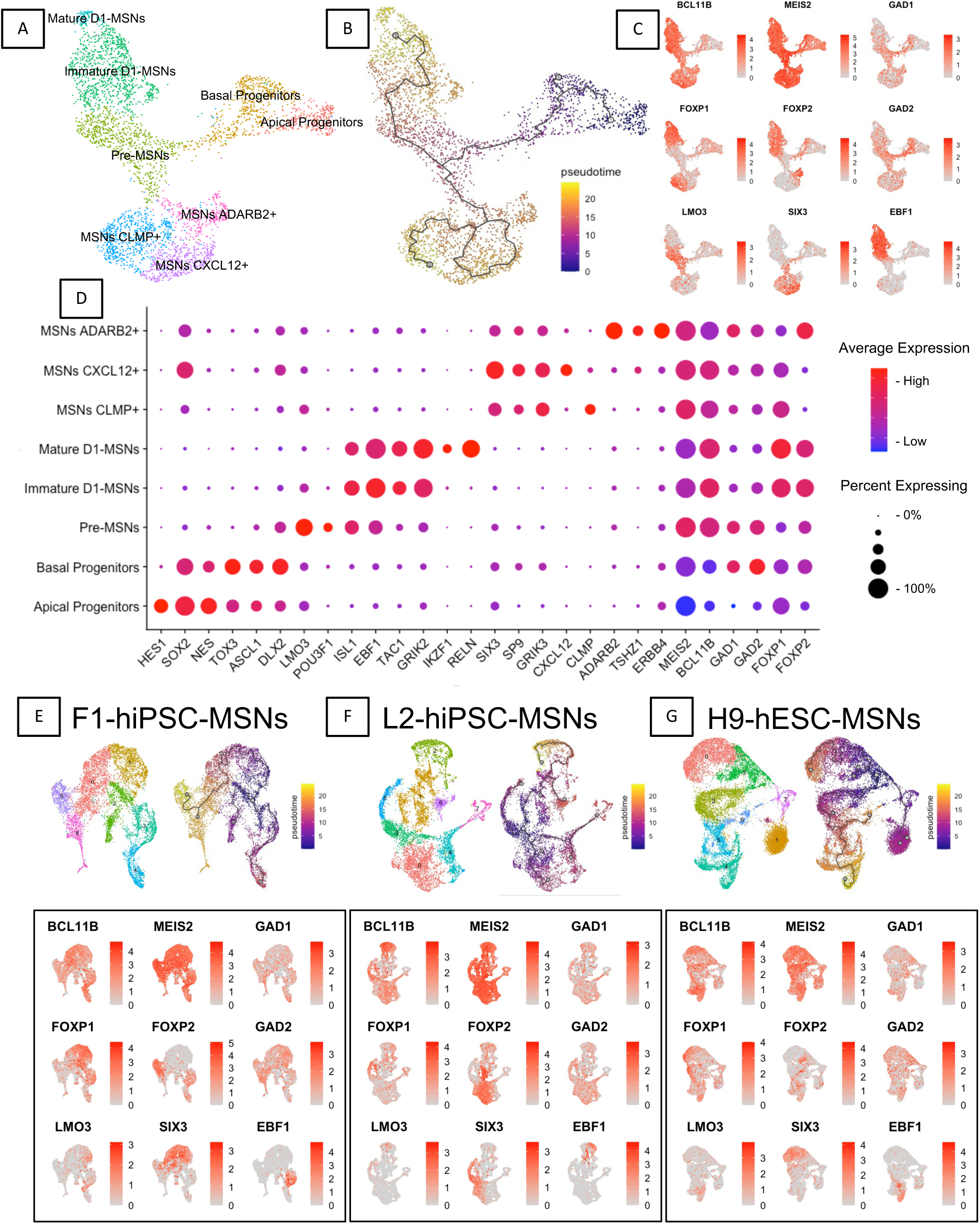
– Individual scRNAseq analysis of MSNs derived from authentic LGE and hPSC-MSNs **A.** UMAP of primary LGE (pLGE) derived cells of MSN lineage, annotated using canonical gene expression. **B.** Pseudotime analysis trajectory plot mapped onto pLGE UMAP depicted in A, showing predicted developmental path from apical progenitors to maturing MSNs with a major fate choice in the pre-MSN stage. **C.** Expression plots of key MSN markers in pLGE. **D.** Dotplot showing expression of key MSN markers across pLGE clusters, coloured according to average expression across a cluster’s population and percentage of cluster expressing the gene. **E-G.** Summary UMAP plots showing clusters, pseudotime trajectory, and expression of key MSN markers in hiPSC-MSNs, expanded in Figures S6-8.

Similar analysis was conducted on each population of hPSC-MSNs (Figure 2 E-G, Figure S6-8). Overall we observed an enriched MSN phenotype in our hPSC-MSNs, exhibiting similar patterns of expression as in the primary LGE MSNs. For instance, global MSN markers BCL11B and MEIS2 were enriched across the populations, as were GABAergic marker genes GAD1 and GAD2, indicating a strong commitment to an MSN-like phenotype. Markers consistent with early progeny and pre-MSNs (e.g. HES1, TOX3, NES, ASCL1, DLX2 and LMO3) were present within the populations even at this late time point, but only in a low proportion of cells, indicating the majority had committed to a more mature phenotype. Across all three hPSC-MSN populations we observed enrichment of MSN subtype genes (e.g. EBF1, TAC1, SIX3, SP9), and markers of maturing phenotypes (e.g. GRIK2, RELN, IKZF1, GRIK3, CLMP, CXCL12, ADARB2), indicating hPSC-MSNs had successfully differentiated to the similar subtypes as authentic LGE derived MSNs. However, while D1 subtypes (BCL11B/MESI2/EBF1+ cells) were present, they were underrepresented in our differentiations. Conversely D1/D2 phenotypes (BCL11B/MEIS2/ADARB2+ MSNs) made up a larger proportion of the final cell population when compared to primary LGE derived MSNs, particularly in the L-iPSC-MSNs.

### Despite exhibiting canonical MSN genes, there is evidence that hPSC-MSNs are not equivalent to authentic LGE derived MSNs

Having established that hPSC-MSNs exhibited similar cell types and signatures to authentic counterparts, we next began direct comparison with authentic tissues. We projected the hPSC-MSN cells onto the fetal LGE UMAP, to assign cluster identities using map-based expression similarities (Figure 3 A). Similarity scores were then created for each cell within a given cluster, against the average expression for that cluster within the fetal LGE sample (Figure 3 B). hPSC-MSNs consistently exhibited strong similarity to authentic tissues, with average expression of sub-type genes being expressed at comparable levels. However, for each sub-type of hPSC-MSN cell, there were sub-populations with lower similarity scores, indicating a divergence from the authentic tissue. This was most apparent in the mature D1 and D2 phoenotypes, which may indicate that at least some of this variation is driven by an increase in maturity between hPSC-MSNs and MSNs collected from the fetal brain.

**Figure 3.**
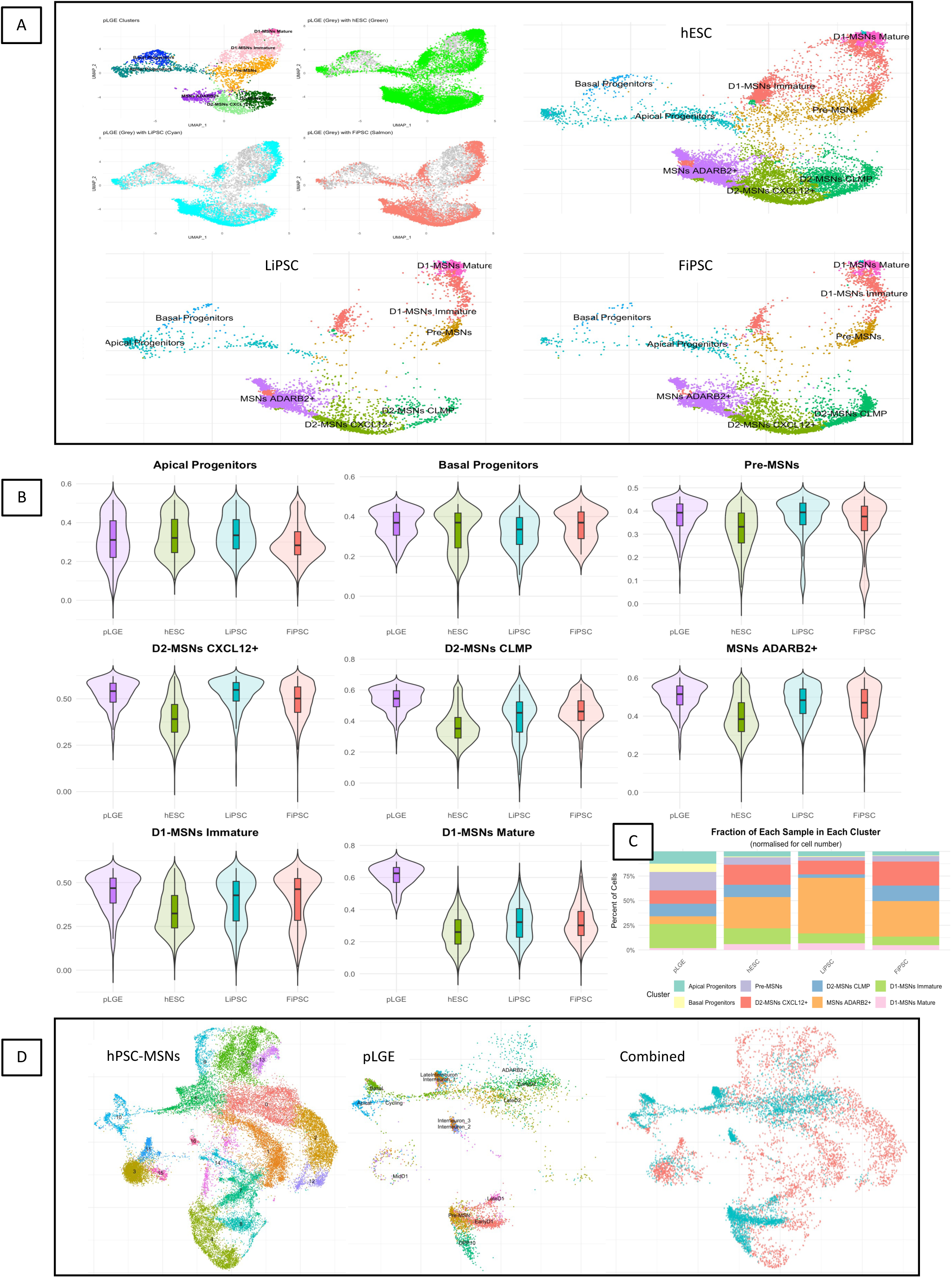
– Projection based scRNAseq analysis of MSNs derived from authentic LGE and hPSC-MSNs **A.** UMAPs of hPSC-MSNs projected onto LGE, exhibited both with (upper left small) and without (larger) LGE sample. **B.** Box plot with violin density overlay exhibiting similarity index scores for each sample cell subtype, measured against the average similarity score for authentic LGE subtypes, greater value indicates greater similarity where 1 = complete similarity. **C.** Percentage of each population, broken down by cell sub-type. **D.** UMAPs of LGE cells projected onto integrated hPSC-MSNs, showing integrated hPSC-MSNs alone (left), LGE alone (center) and combined (right).

Additionally, hPSC-MSNs exhibited fewer total cells in immature cell subtypes and greater numbers of cells in mature populations (Figure 3 C). Reverse projection mapping (projecting fetal LGE onto hPSC-MSNs) further evidenced this possibility, as fetal LGE tissues did not cluster at further edges of the hPSC-MSN clusters (Figure 3 D), which exhibited maturity markers such as RBFOX3. However, the least mature cells in hPSC-MSNs samples (apical progenitors) exhibited a greater number of immature markers, suggesting that the divergence in this population was driven by cells that were failing to exit early cell cycling and remaining in a proliferative state.

Of the hPSC-MSNs, LiPSC-derived MSNs exhibited the highest similarity scores for apical progenitors, pre-MSNs, CXCL12+ D2 MSNs, CLMP+ D2 MSNs, ADARB2+ MSNs, and Mature D1 MSNs. Conversely, FiPSC derived MSNs held the highest average similarity for Early D1 MSNs, and hESCs hald the highest average similarity for basal progenitors. Overall this supports the concept that the primed LGE derived hiPSCs maintain sufficient epigenetic memory to influence their differentiation into more authentic MSNs when compared to naïve hESCs or non-primed hiPSCs.

A critical limitation of projection mapping is an inherent bias to favour similarities between populations, which may lead to overlooking significant differences. To address this, we next transitioned to an integrated analysis, providing a second method to probe the data (Figure 4). As before, there was considerable overlap between hPSC-MSNs and fetal LGE, corroborating initial assessments (Figure 4 A-D). Yet, there were some notable exceptions, specifically we observed several cell populations (clusters 2, 3, 4, 10, 11, & 16) to be primarily comprised of hPSC-MSNs indicaing they were not represented in the authentic tissue (Figure 4D). Both clusters 2 and 3 expressed high levels of CLMP, potentially indicating these are CLMP expressing D2 MSNs. However only cluster 3 expressed established D2 genes SIX3 and SP9, indicating that cluster 2 may possess an incomplete or abberant phenotype. Comparisons of cluster 3 (CLMP/SIX3/SP9+ but containing ∼0% Primary LGE cells) to cluster 15 (which also contained CLMP/SIX3/SP9+ cells and contained ∼59% primary LGE cells) revealed 292 differentially expressed genes: cells in cluster 3 expressed genes associated with neuronal projections and axon regulation, whereas cells in cluster 15 expressed genes associated with fetal brain tissues, indicating that this division is likely due to differences in maturity between these populations. Similarly, cells in cluster 10 expressed D1 marker EBF1, but did not express genes that differentiated between immature and mature D1 cells in the primary LGE tissues (e.g. ISL1 or RELN), potentially indicating that these cells are an incomplete D1 phenotype as they have exited the early ISL1+ phase but not yet begun exhibiting typical signs of maturity. Whereas clusters 4, 11, and 16 contained cells positive for D1/D2 genes such as ADARB2, which only represented a small proportion of the overall population in primary LGE, likely emphasising a bias for our protocol to over-produce the D1/D2 phenotype (Figure 4 F).

**Figure 4.**
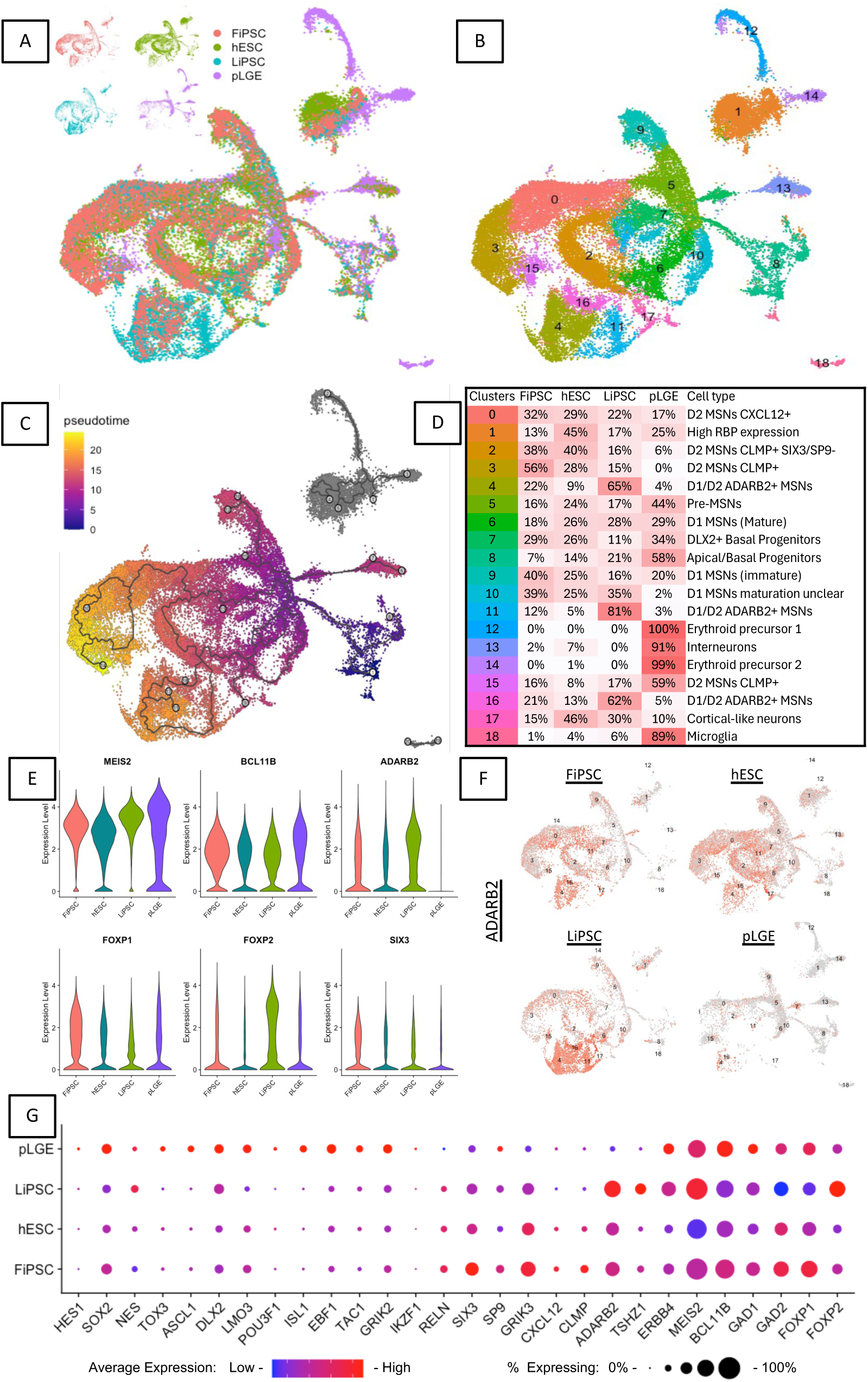
– Integrated scRNAseq analysis of MSNs derived from authentic LGE and hPSC-MSNs **A.** UMAPs of integrated data set, showing overlap between cell populations (main), and individual maps of each sample (small). **B.** Cell clusters depicted on UMAP plot. **C.** Pseudotime analysis trajectory plot mapped onto UMAP. **D.** Table depicting each cluster’s predicted phenotype, and the contributing proportion by sample. **E.** Violin plots showing differences in expression level across total populations for differentially expressed striatal genes. **F.** Expression plots for ADARB2 split by sample **G.** Dotplot showing expression of key MSN markers across samples, coloured according to average expression across a sample’s population and sized by percentage of cells expressing the gene.

### Line specific differences between hPSCs heavily impact on final cell population compositions, but not subtype phenotype

Lastly, we compared the hPSC-MSNs produced from each cell line to explore how impactful the specific hPSC line is on final population composition. Between L2-and F1-hiPSC-MSNs, there were 89 genes expressed at significantly different levels, 31 higher in the F1-hiPSC-MSNs and 58 higher in the L2-hiPSC-MSNs. These differences primarily reflected a difference in subtype proportions, for example, greater levels of FOXP1, ZNF503, SIX3, CLMP, and GRIK3 expression were observed across the F1-hiPSC-MSNs, indicating higher yields of both D1-like and D2-like MSN subtypes than in the L2-hiPSC-MSNs; conversely, greater proportions of ADARB2, TSHZ1, FOXP2, ERBB4, and LUZP2 expressing cells were observed in L2-hiPSC-MSNs, indicating a higher yield of D1/D2-like MSNs than in the F1-hiPSC-MSNs (Figure 4 E-G).

Q-PCR analysis exploring the relative expression a pannel of MSN related genes were used to confirm these differences were consistent between lines and across multiple differentiations. As in our scRNAseq analysis, we observed F1 cultures consistently produced populations exhibiting higher expression levels of FOXP1 (vs all: F_8,20_=5.82, *p=0.001*; Figure S9), ZNF503 (vs L11-, but not L2-: F_4,10_=3.54, *p=0.048*; Figure S9), DRD2 (vs all, F_8,20_=7.17, *p<0.001*) BCL11B (vs L2 and L11, F_4,10_=12.15, *p=0.001*) and DARPP32 (vs all, F_8,20_=11.48, *p<0.001*; Figure S9).

Further exploration of specific MSN subtypes did not reveal gene expression profiles unique to individual lines, indicating that the epigenomic background of our hiPSC lines mainly effected their potential to yield total MSNs and subtype fate commitment. Specifically this caused the F1 line to yield greater proportions of D1-and D2-like MSNs, and the L2 line to yield greater numbers of ADARB2+ D1/D2-like MSNs, in spite of their isogenic nature.

## Discussion

Human pluripotent stem cells (hPSCs) are a powerful research tool due to their accessibility and capacity to differentiate into virtually any cell phenotype. However, despite strenuous efforts to optimise differentiation protocols, it is becoming increasingly clear that hPSC derived cells are not always equivalent to their target phenotype. Additionally, variability exists between type of hPSC (i.e. induced vs embryonic), between individual cell lines, and even across passage number (Ortmann and Vallier, 2017). Understanding the extent, origin, and implications of this variability is essential for ensuring the fidelity of hPSC-based models. Here, we asked how epigenetic variability can influence the differentiation capacity of isogenic hiPSCs toward a medium spiny neuron (MSN)-like phenotype.

Our data reveal that subtle but consistent epigenetic signatures persist through hiPSC reprogramming and subsequent neuronal differentiation. This finding is consistent with previous studies reporting similar retention of tissue-specific methylation in iPSCs derived from neural tissues (Tian et al., 2011; Hargus et al., 2014; Roost et al., 2017). These stable signatures were associated with differential DNA methylation, altered gene expression patterns and consistent preferences in subtype yields in the resulting hPSC-MSNs. Since these lines are isogenic, the most plausible driver of these differences is epigenomic variability, including inherited memory from the tissue of origin.

While LGE-derived hiPSCs displayed some beneficial epigenetic features, such as reduced hypomethylation of off-target pathways and enhanced expression of genes associated with the target cells, this did not greatly enhance MSN fidelity. In fact, despite being epigenetically non-primed, fibroblast-derived hiPSCs were superior than LGE-derived hiPSCs by some metrics, such as global CTIP2 and DARPP32 expression. One possible explanation for this is that the differentiation protocol, was originally optimised for naïve hESCs (Arber et al., 2015) and may not be suited to the epigenetic context of neural-primed cells. Thus, pathway signalling conditions (such as timing, dosage, etc) that favour naïve lines may prove suboptimal or disruptive when applied to cells with a pre-existing striatal bias. This possibility underscores the complex interplay between intrinsic cellular epigenetics and extrinsic protocol design.

DNA methylation plays a key role in regulating gene expression and cellular identity. During iPSC reprogramming, widespread changes in DNA methylation occur, leading to the loss of the majority of tissue-specific regulatory landscapes and the acquisition of a more naïve methylomic state (Papp and Plath, 2013). Here, we observed a pronounced increase in methylation in LGE-derived hiPSCs relative to the original fetal LGE tissues, with many of the affected genes associated with striatal function. Upon differentiation to hPSC-MSNs, we observed partial reversal of these methylation changes, particularly at genes commonly used to assess striatal identity such as BCL11B, MEIS2, FOXP1 & FOXP2. However, the methylomes of all hPSC-MSNs remained substantially different from those of authentic fetal MSNs, with large numbers of genes associated with striatal function remaining hypermethylated. This was unexpected, as the fetal LGE tissue used as a reference is inherently heterogeneous, comprising cells across multiple developmental stages and lineages (Bocchi et al., 2021; Shi et al., 2021). Considering the binary nature of DNA methylation, one would expect increased heterogeneity within the LGE tissues to reduce the tissue specific signature, and for a phenotype-focused population (as hPSC-MSNs are supposed to be) to therefore exhibit a stronger signature if they had successfully acquired it. Despite this, the LGE tissues consistently exhibited greater demethylation across striatal, MSN, and general neuronal genes. This suggests that while current protocols largely induce relevant epigenetic programs, they fail to fully restore the authentic MSN methylome. Interestingly, the genes which exhibited successful changes included markers commonly used to measure the success of a given MSN protocol. When these genes are the metrics by which a differentiation protocol is deemed successful, is it really surprising that lesser studied genes and functions are left behind?

Interpreting the consequences of these epigenetic differences is challenging, as the role of methylation varies by genomic context and is not uniformly associated with repression or activation (Greenberg & Bourc’his, 2019; Smith & Meissner, 2013). Here, our scRNAseq data show that hPSC-MSNs recapitulate major aspects of authentic MSN development, including transcriptional transitions from progenitors to differentiated subtypes. We identified populations corresponding to apical and basal progenitors, pre-MSN intermediates, and differentiated MSNs, in both fetal and hPSC-derived samples. Similarity indexes suggested that overall these cells were highly comparable. However, differences in relative subtype proportions were evident, and some hPSC-specific clusters lacked complete MSN expression signatures, suggesting incomplete or divergent differentiation. This was particularly evident for both the most immature and most mature cells, which expressed canonical MSN markers but appeared to lack key genes that distinguish authentic subtypes, supporting the notion that epigenetic or protocol-related factors produce cells that mimic, but do not fully match, genuine MSN phenotypes.

Taken together, our findings support three major conclusions. First, hPSC lines differ in their differentiation capacity, even when isogenic. Second, tissue-specific epigenetic memory persists through reprogramming and differentiation, and this residual memory can shape differentiation outcomes (both positively and negatively) depending on protocol context. Third, hPSC-derived cell populations, even when expressing canonical marker genes, may include transcriptionally and epigenetically incomplete or inauthentic subtypes. This highlights the importance of using multimodal and unbiased epigenetic, transcriptomic, and functional validation when evaluating hPSC-based models. Future work may benefit from refining differentiation protocols to better account for the epigenetic starting state of each line, or from directly modifying epigenetic states (e.g. via CRISPR-based editing) to more closely resemble authentic developmental trajectories. As single-cell methylome technologies become more accessible, these tools will offer unprecedented resolution for understanding how epigenetic identity constrains or enables authentic cell fate acquisition.

In summary, while hPSC-derived MSNs broadly resemble their *in vivo* counterparts and successfully model many aspects of MSN development, important epigenetic and transcriptional differences remain. Recognising and addressing these limitations is essential for improving the fidelity of hPSC-based models and for realising their full potential in research and therapeutic development.

## Funding declarations and acknowledgements

This work was funded by Wellcome Trust studentship grants 105215/Z/14/Z and 105215/Z/14/A, MRC grant 518008, and a Cardiff University NMHRI seedcorn grant. The authors would like to acknowledge and thank the technical expertise of the SWIFT team in the Brain Repair Group at Cardiff University and at University Hospital Wales, and the Genomics Research HUB at Cardiff University, Lesley Bates and Joanne Morgan.

## Author Contributions

Conception and design of the study: OJMB, NNV, NMW, SVP and AER. Acquisition of data: OJMB. Analysis and interpretation of data: OJMB. Drafting or revising the manuscript: OJMB, MJL, SVP, and AER. All authors have approved the final article.

## Methods

### Primary Tissue Culture

All primary human fetal tissue was acquired from the South Wales Initiative for Fetal Transplantation (SWIFT) research tissue bank. It was dissected and dissociated as previously described (Roberton et al., 2018).

### Generation of hiPSCs

Primary tissues were cultured on PDL (Gibco) treated nunc cell culture plates (Thermofisher). Neural tissues were maintained in a neural media (DMEM/F12, +2% B27 Supplement 50x (B27; Gibco), +1% fetal bovine serum (FBS; Gibco), +1% Penicillin-Streptomycin (PS; Gibco)), and fibroblasts were cultured in a basic media (DMEM/F12, +10% FBS, +1% MEM Non-Essential Amino Acids Solution 100x (MEM-NEAA, Gibco), +1% L-Glutamine (Gibco), +1% PS). 3 days following dissociation and plate down we induced pluripotency via the CytoTune -iPS 2.0 Sendai Reprogramming Kit (Invitrogen), at MOI = 5 for 24 hours. Tissues were maintained under pluripotent culture conditions from day 5 post-infection.

### Culture of hPSCs

hPSCs (hiPSCs and hESC line H9 (WA09, Wicell)) were maintained on Geltrex (Gibco) or hESC-qualified Matrigel (Corning) coated nunc plates, in E8 or E8-Flex media (Gibco). hPSCs were passaged when between 70-90% confluency using EDTA (Sigma) or ReLeSR (StemCell Technologies).

### Spontaneous differentiation of hPSCs

hiPSCs at 40% confluency were cultured in KSR medium (KnockOut DMEM (Gibco), +10% KnockOut Serum Replacement (Gibco), +1% L-glutamine, +1% MEM-NEAA, +1% PS, +0.1% β-mercaptoethanol (Gibco)) for 5 days to allow for spontaneous differentiation.

### Directed differentiation of hPSCs towards MSN-like fate

hPSCs were differentiated towards an MSN-like fate using a protocol adapted from previous works (Arber et al., 2015, Repair HD; Figure 1A).

Prior to differentiation, hPSCs were cultured on Growth Factor Reduced Matrigel (Corning; diluted 1:15 in DMEM-F12 and coated for 1 hour at 37°C) in E8 or E8-Flex media until ∼70% confluency (day 0). Differentiation was initiated by changing media to N2B27-RA media (2:1 ratio of DMEM/F12 and Neurobasal medium (Gibco), +1% L-glutamine, +0.6% N2 supplement 100x (N2; Gibco), +0.6% B27 without retinoic acid supplement 50x, (B27-RA; Gibco), +0.2% MycoZap plus CL (Lonza), +0.1% β-mercaptoethanol), supplemented with 10 μM SB-431542 (SB; Tocris), 200 nM dorsomorphin (R&D),100 nM LDN-193189 (LDN; Tocris) and 2 μM XAV-939 (Xav; Tocris). From day 6, SB was removed from media. On day 7, cultures were passaged using EDTA at a 2:3 ratio split onto PDL + Fibronectin (Millipore) coated nunc plates. From day 10, dorsomorphin and LDN were removed from media, and 25 ng/ml Activin A (R&D) was added. On day 16, cultures were passaged using Accutase (Sigma) and frozen into N2B27-RA media supplemented with 10% DMSO (Sigma) and stored at -80°C. Following thawing, cells were plated at ∼100k cells/cm^2^ on PDL+Laminin (Sigma) coated nunc plates in N2B27-RA media supplemented with Xav and Activin. From day 18, Xav and Activin were removed from media. From day 20, media was replaced with N2B27 (as N2B27-RA, but B27-RA is replaced with B27), supplemented with 10 ng/ml BDNF (Peprotech) and 10ng/ml GDNF (Peprotech). Medium was replaced every second (N2B27-RA) or third (N2B27) day throughout. Cells were maintained up to day 40.

### Fluorescent immunocytochemical staining and analysis

Cells were fixed in 3.75% PFA (Millipore) and permeabilised in 100% molecular grade ethanol (Sigma). Fluorescent immunocytochemistry was performed using standard protocols, including incubations in blocking solution, primary antibodies, and secondary antibodies.

Primary antibodies: α-FP (Abcam), α-SMA (Sigma), βIII-tubulin (Sigma), CTIP2 (Abcam), DARPP32 (Abcam),DESMIN (Abcam), MAPa/b (Sigma), NANOG (Abcam), NESTIN (Neuromics), OCT4 (Abcam), SOX2, (Abcam), SSEA3/4 (Abcam), TRA-1-60 (Abcam), Vimentin (Millipore).

Stained cells were visualised using either a ZEISS Axio Imager 2 upright fluorescent microscope with Axiovision software, or using a Leica DMI6000B inverted fluorescent microscope with LAS X software. Cell counts were conducted using 5 images per sample and three samples per condition. Statistical analysis was conducted using SPSS (IBM).

### Gene expression analysis using PCR and qPCR

RNA was extracted using RNeasy Mini-Prep (Qiagen). cDNA was synthesised using SuperScript IV First-Strand Synthesis System (Thermofisher). PCR was conducted using BioTaq DNA polymerase (Bioline). qPCR was conducted using PowerUp SYBR Green (Invitrogen). For qPCR, data is expressed as ΔΔCt, normalised to the average Ct of two reference genes (GAPDH and β-ACTIN), and to the gene expression in H9-MSNs at day 30. Fold change expression values were used for statistical analysis. Statistical analysis was conducted using SPSS (IBM).

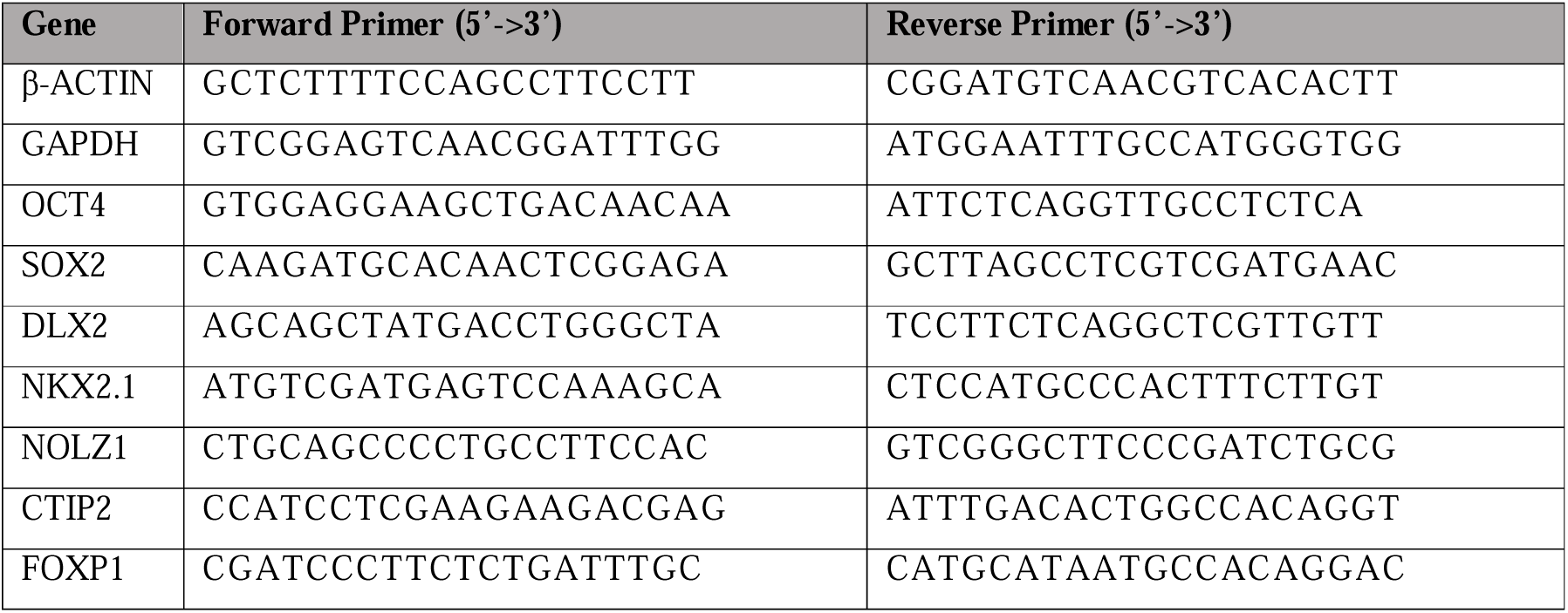
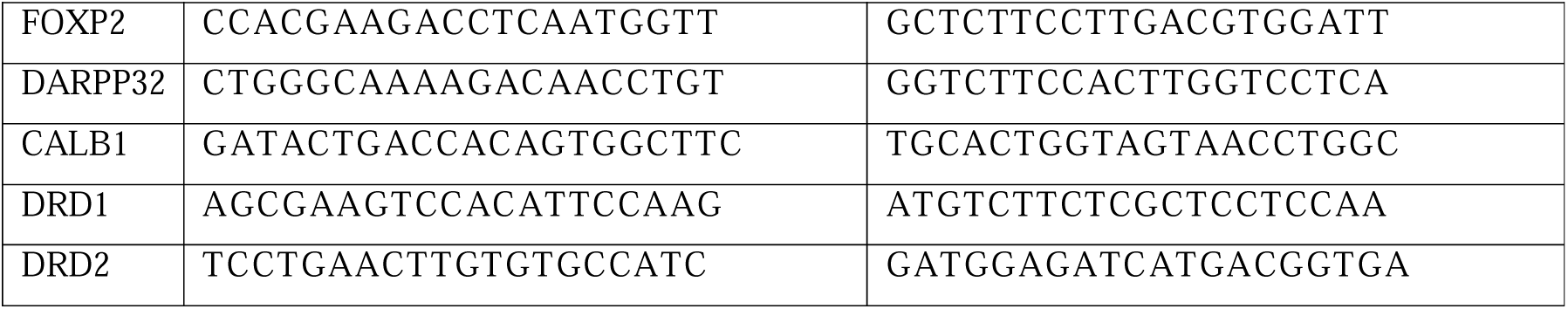

### DNA Methylation Analysis

DNA methylation analysis was conducted as previously described (Choompoo et al., 2021; Morris et al., 2014; Aryee et al., 2014; Teschendorff et al., 2012).

### scRNAseq Analysis

scRNAseq performed using the Chromium Single Cell 30 v2 platform (10X Genomics). For each sample ∼10,000 viable cells were collected for library generation. Libraries were sequenced with the Illumina NovaSeq platform, using paired end sequencing with single indexing. Raw data files (Fastq) were demultiplexed, processed, aligned and quantified using the Cell Ranger Single-Cell Software Suite (10x Genomics).

Downstream analysis was performed using Seurat5 (Satija et al., 2015; Butler et al., 2018; Stuart et al., 2019; Hao et al., 2021; Hao et al., 2024). Cells with <200 or >7500 genes were removed, as were those with >10% mitochondrial gene expression. For analysis of primary LGE alone, cell clusters expressing non-MSN lineage markers were removed. Pseudotime analysis was conducted using Monocle3 (Trapnell et all., 2014; Qiu et al., 2017; Cao et al., 2019).

### Gene enrichment analysis

Gene sets which were found to be differentially methylated (DNA methylation) or differentially expressed (scRNAseq) were analysed for significant terms using Enrichr (Chen et al., 2013; Kuleshov et al., 2016; Xie et al., 2021). f

---

**Figure S1:**
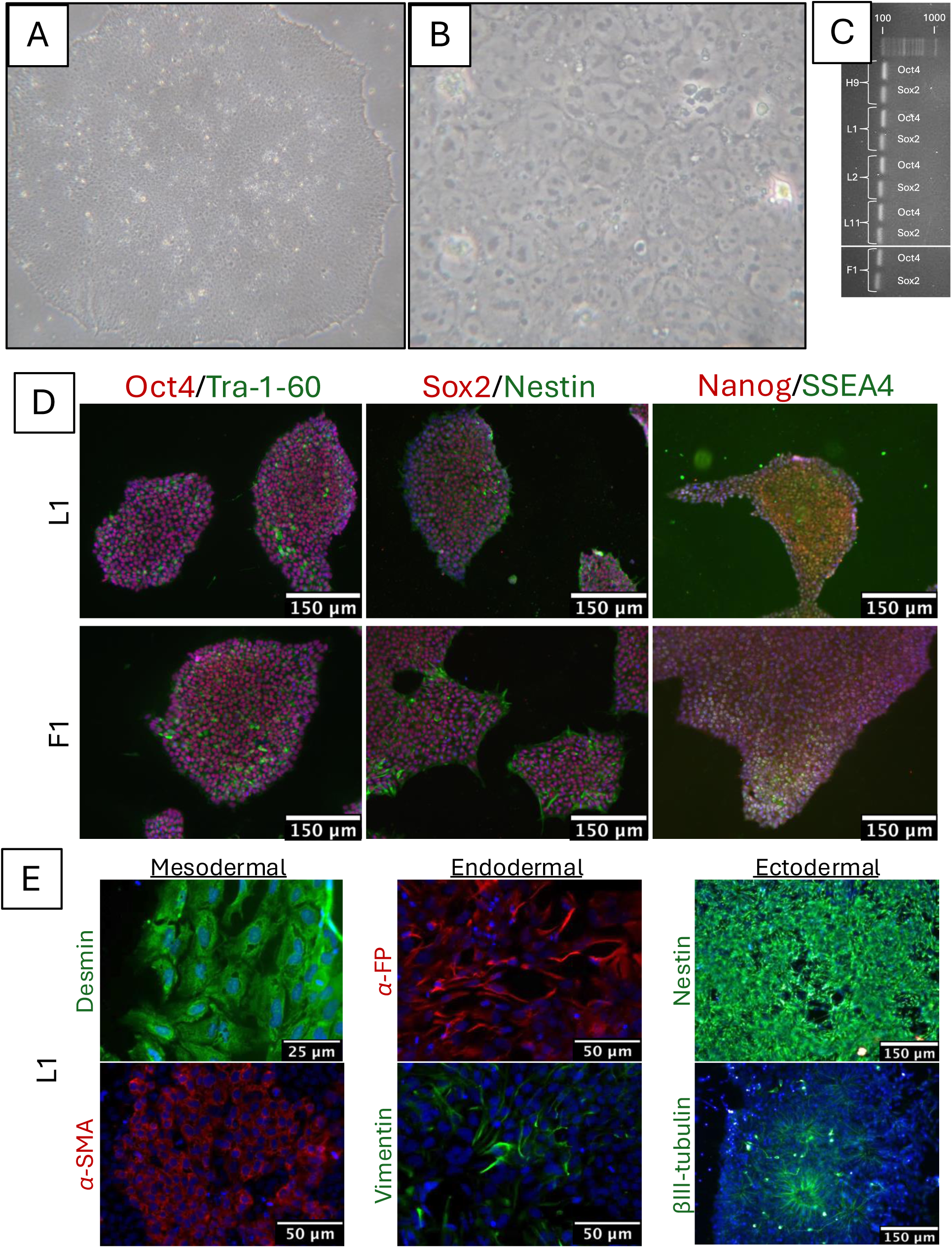
Generation and validation of isogenic hiPSC lines. **A+B.** hiPSCs exhibit typical PSC morphology. Representative brightfield images showing hiPSC colonies were rounded and contained tightly packed cells, which exhibited flat rounded shape with high nucleus to cytoplasm ratio. **C+D.** hiPSCs express classical pluripotent markers OCT4, SOX2, TRA-1-60, NANOG and SSEA4. PCR product visualised following 30 amplification cycles using gel electrophoresis. Representative images of fluorescent-immunocytochemistry. **E.** hiPSCs can produce progeny of all three germ layers following spontaneous or directed differentiation. Representative images of fluorescent-immunocytochemistry staining for markers indicative of each germ layer.

**Figure S2:**
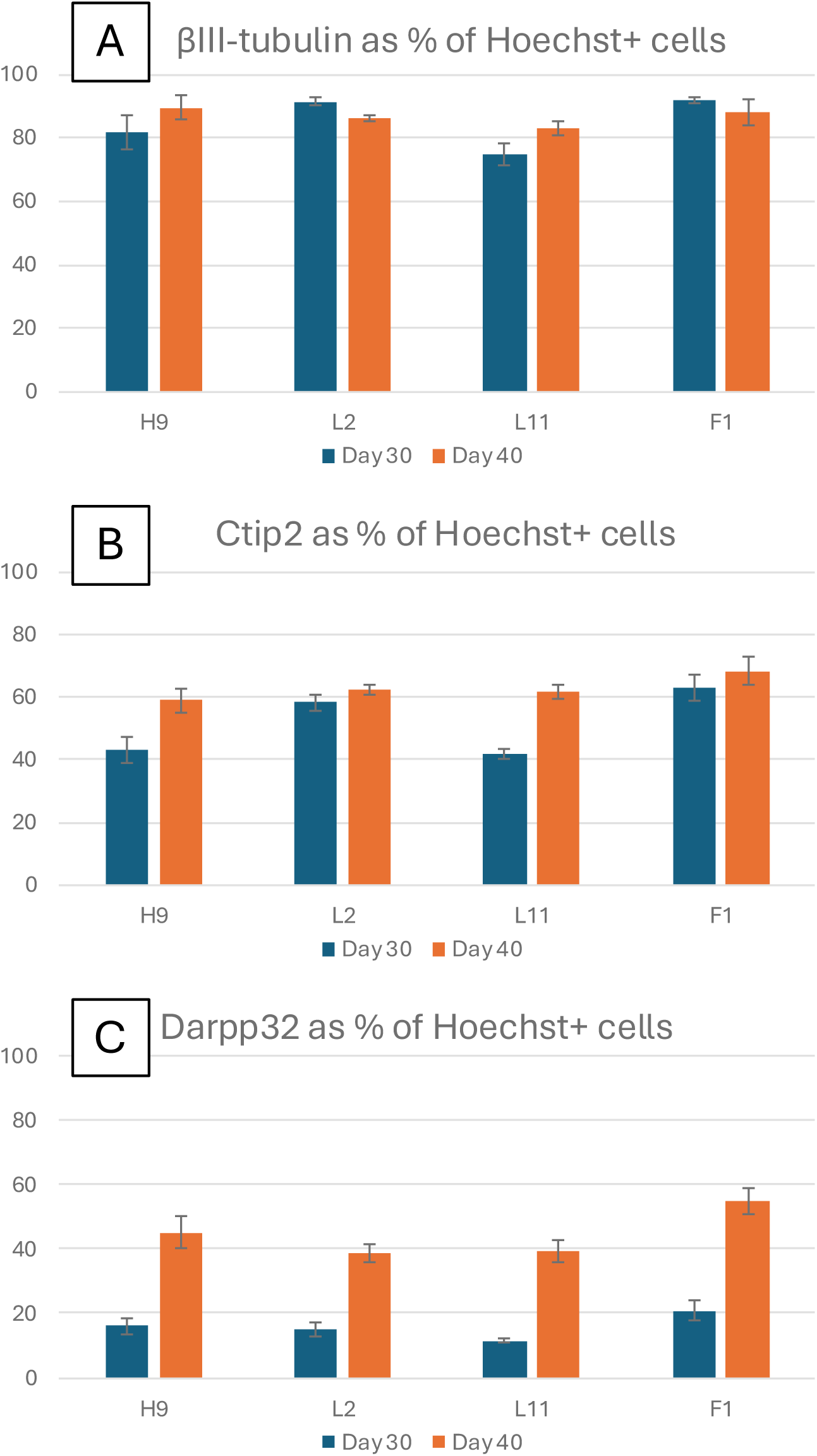
Quantification of immunocytochemical staining. **A-C.** Bar charts depicting the percentage of cells positive for (A) βIII-tubulin; (B) CTIP2/BCL11B; and (C) DARPP32/PPP1R1B. For each line examined, data was collected from 3 independent hPSC-MSN cultures at day 30 and 40 of *in vitro* differentiation.

**Figure S3:**
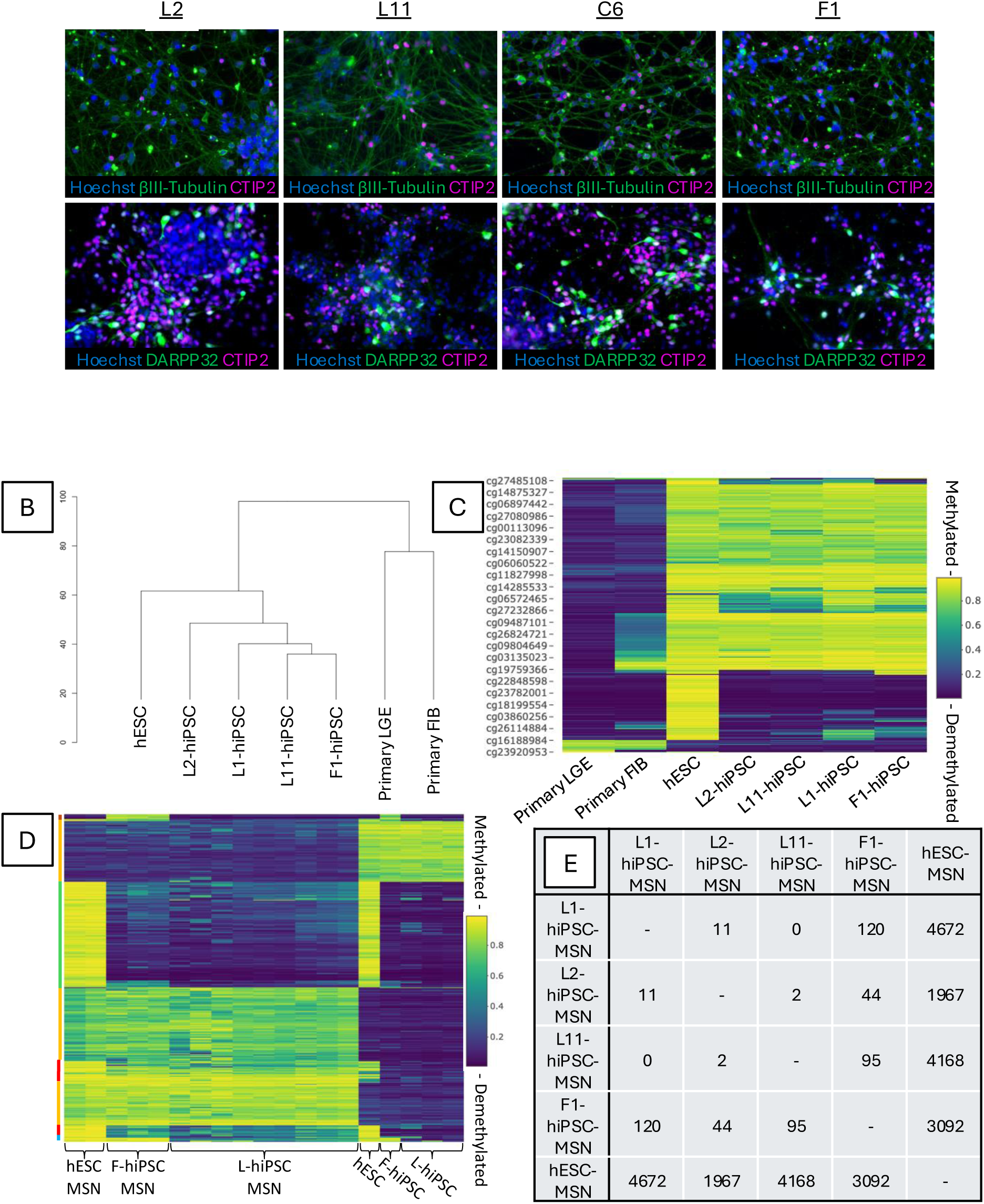
Supplementary DNA methylation images. **A.** hiPSCs Immunocytochemical staining of hPSC-MSNs used in bulk DNA methylation analysis. Cells (Hoechst, blue) are positive for neuronal marker MAP2 (green), and MSN marker DARPP32 (red). **B.** Hierarchical clustering of pluripotent hiPSCs, the original primary tissues from which they were derived, and genomically unique pluripotent hESCs. hiPSCs cluster to hESCs before clustering to tissues from which they were derived. **C.** Heatmap of 1000 most variably methylated probes for analysis conducted in B. **D.** Heatmap of 1000 most variably methylated probes between hPSC and hPSC-MSNs. **E.** Table of differentially methylated probes between hPSC-MSNs, emphasizing fewest differences are found between LGE derived hPSC-MSNs.

**Figure S4:**
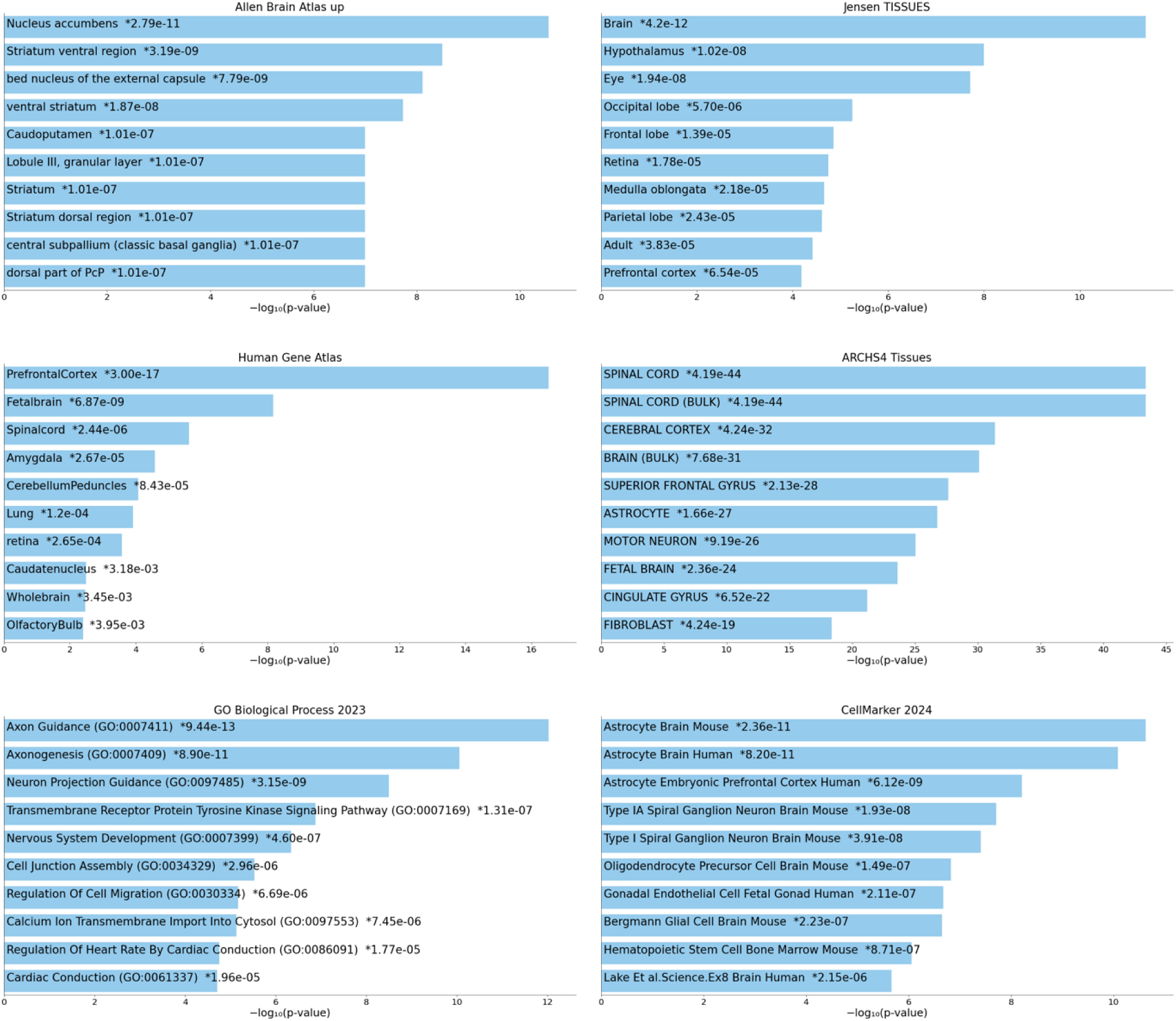
Gene enrichment analysis conducted on differentially methylated genes between Primary LGE and hPSC-MSNs (2224 genes). Bar charts depicting top enriched terms from respective gene set libraries. Based on the -log10(p-value), with p-value next to each term, in order of significance.

**Figure S5:**
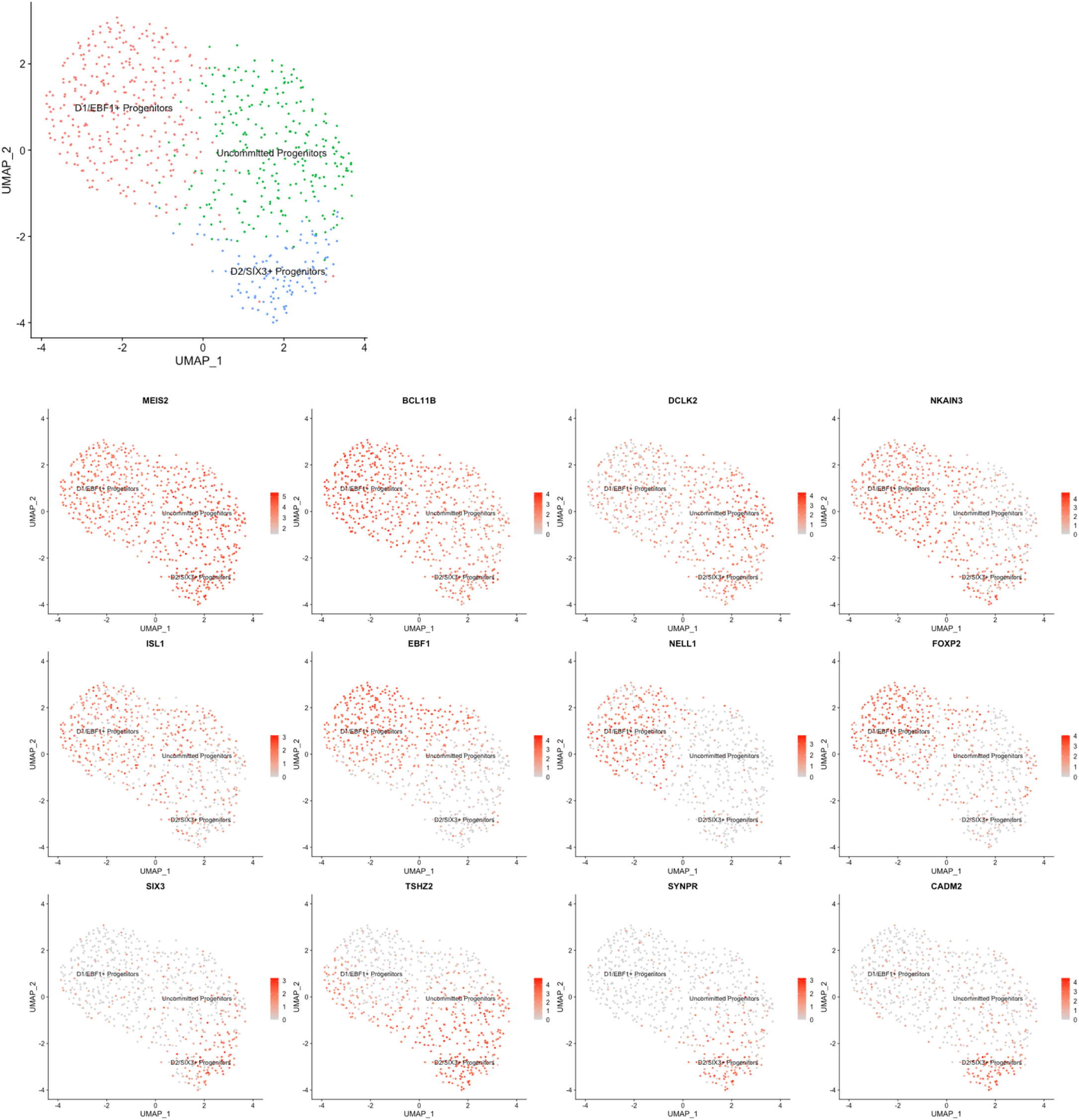
Subclustering of pre-MSN population identified in primary LGE scRNAseq analysis. **A.** UMAP showing sub-clusters within Pre-MSN populations. **B.** Gene expression of canonical MSN genes, D1 and D2 subtype genes, and other upregulated genes.

**Figure S6:**
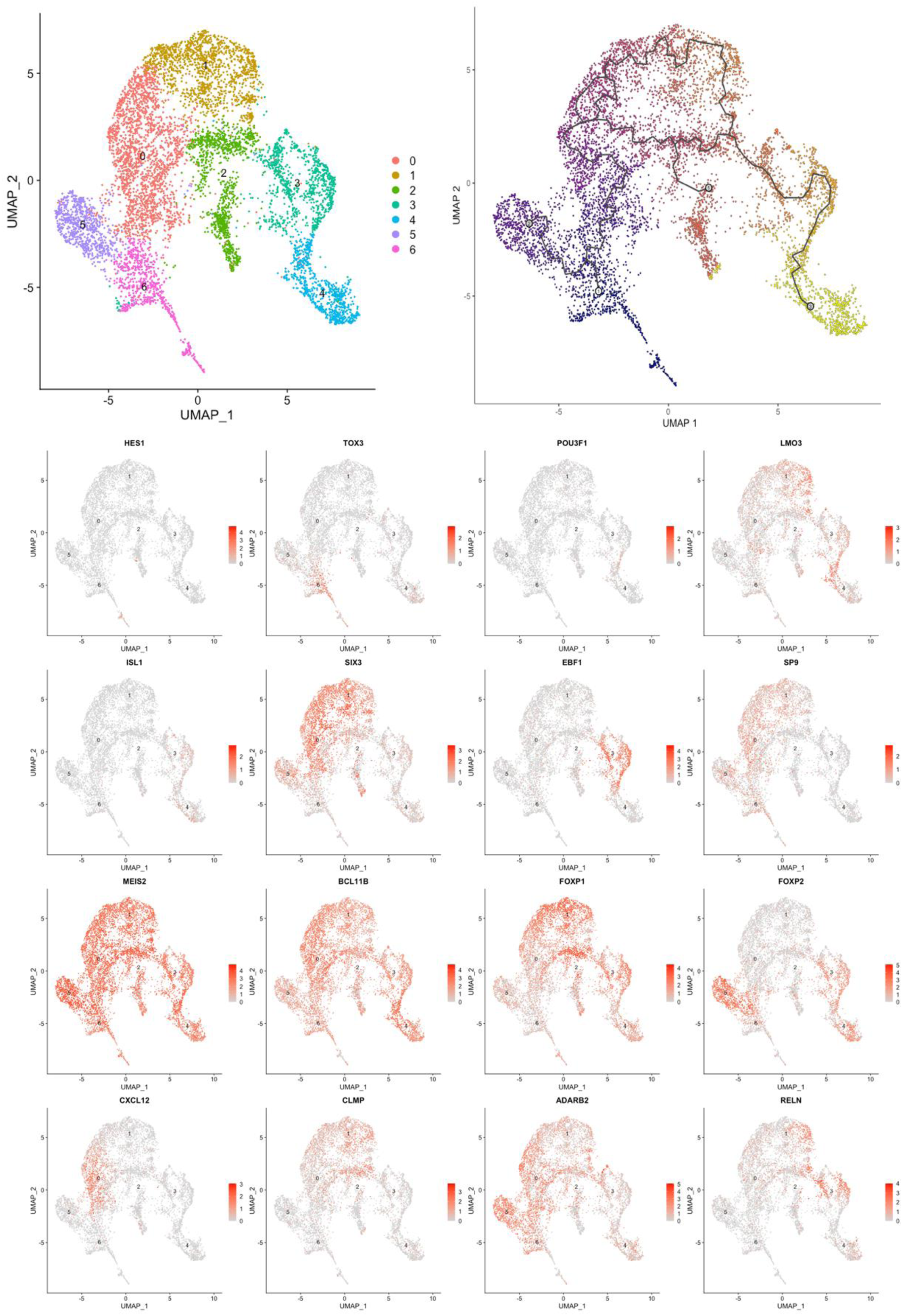
scRNAseq analysis of hPSC-MSNs derived from hiPSC line F1 **A.** UMAP showing clusters identified in population. **B.** Pseudotime analysis. **C.** Gene expression plots of stage defining MSN genes.

**Figure S7:**
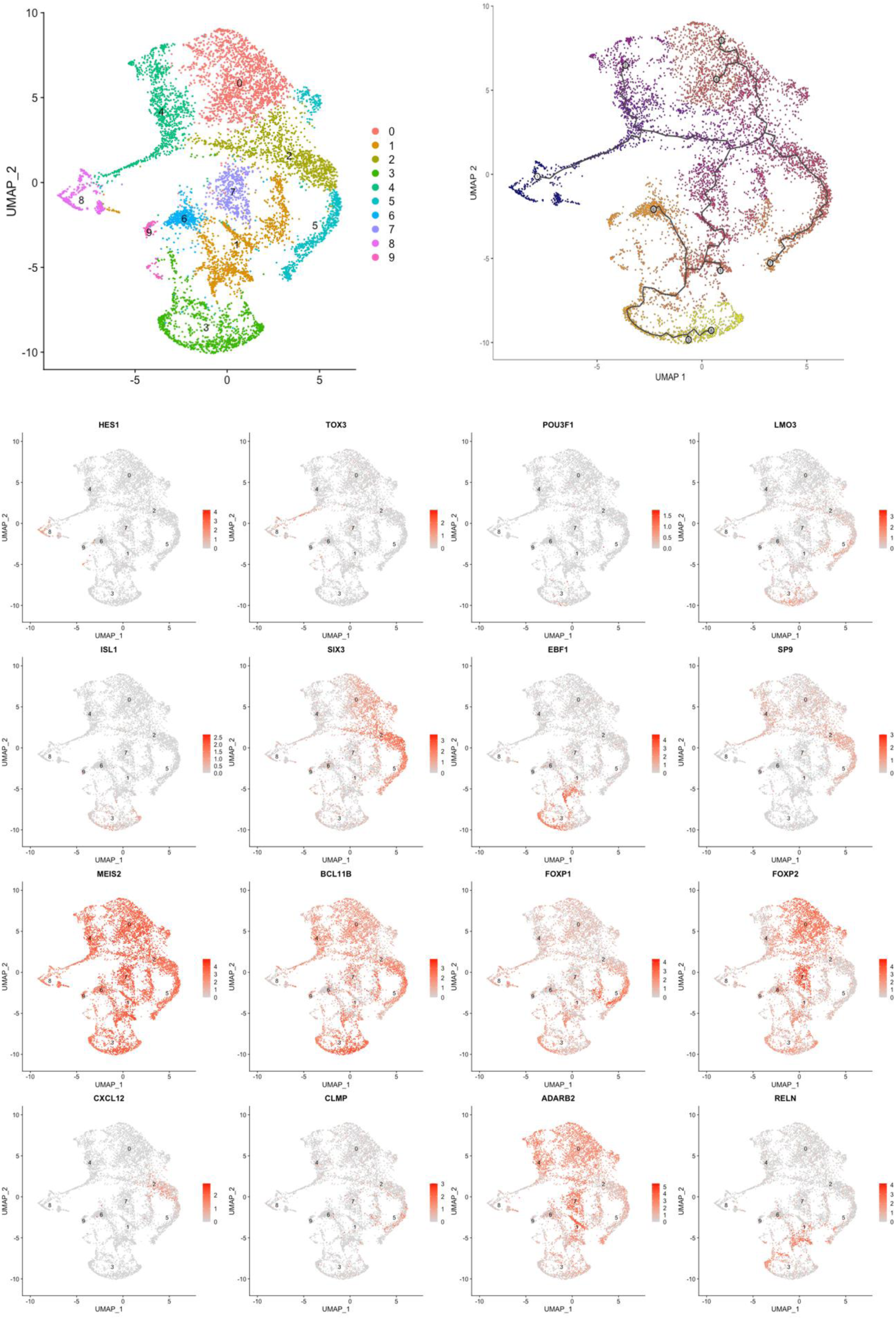
scRNAseq analysis of hPSC-MSNs derived from hiPSC line L2 **A.** UMAP showing clusters identified in population. **B.** Pseudotime analysis. **C.** Gene expression plots of stage defining MSN genes.

**Figure S8:**
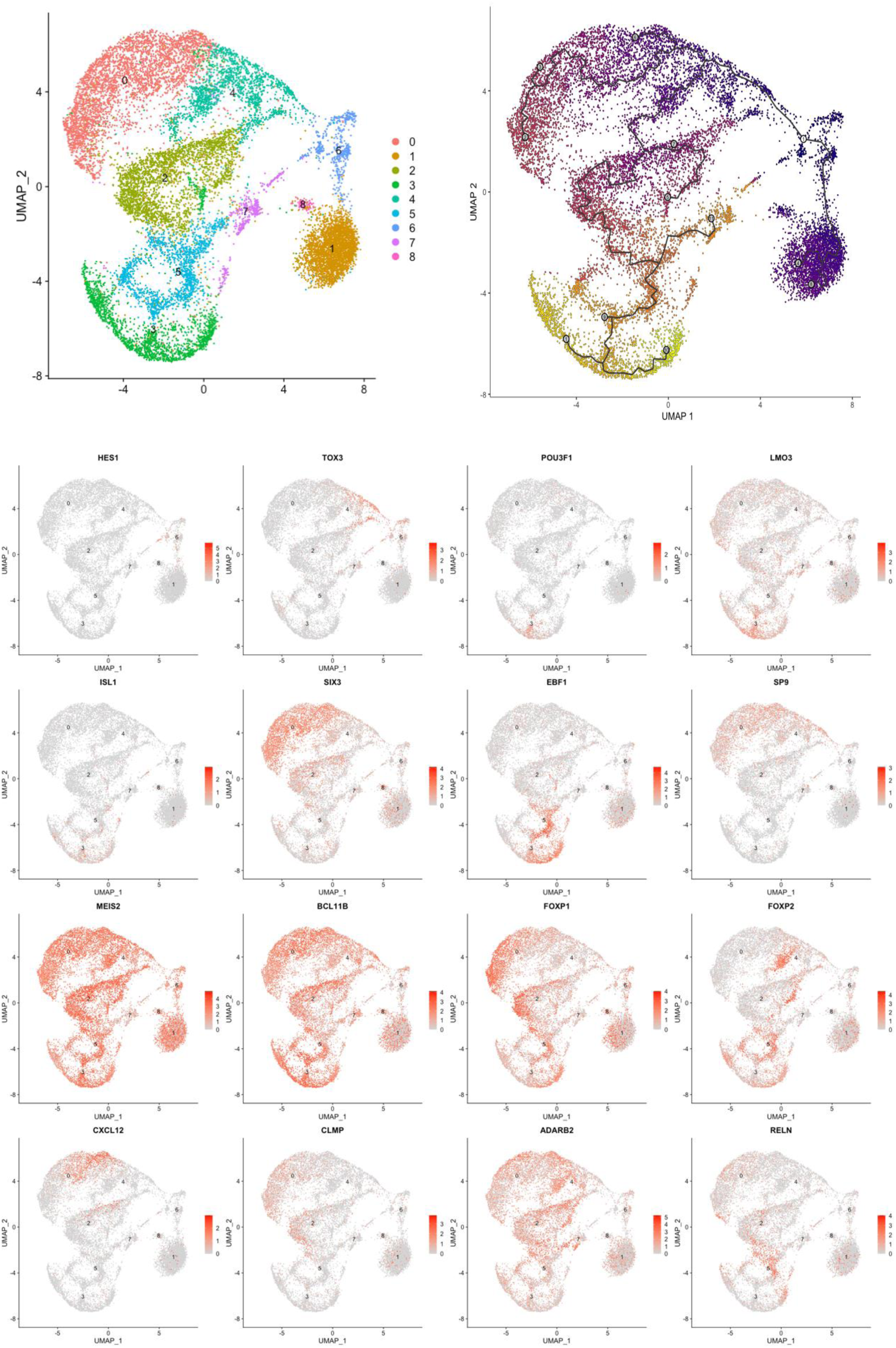
scRNAseq analysis of hPSC-MSNs derived from hESC line H9 **A.** UMAP showing clusters identified in population. **B.** Pseudotime analysis. **C.** Gene expression plots of stage defining MSN genes.

**Figure S9:**
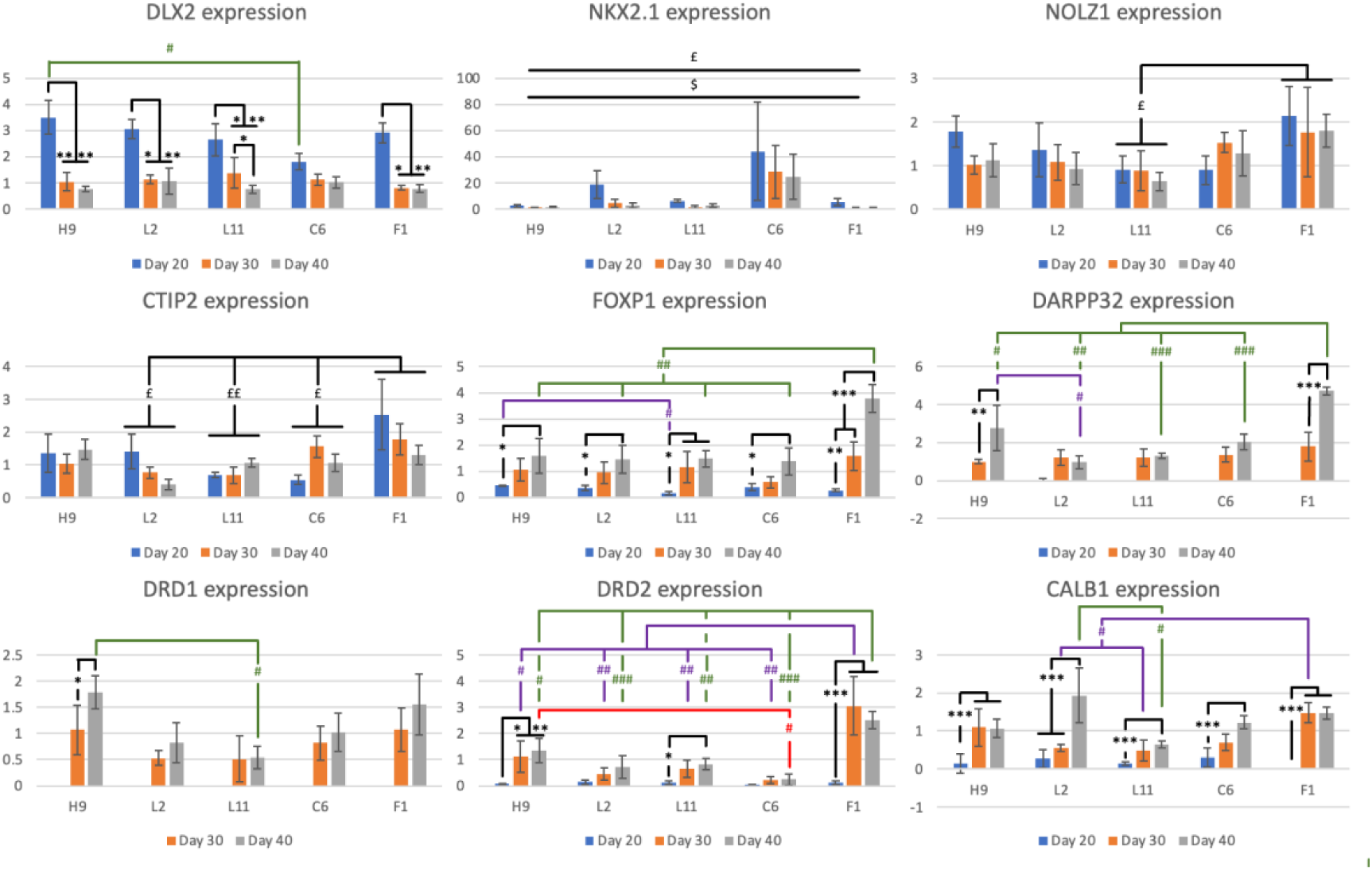
Gene expression analysis using qPCR on RNA collected from L2, L11, C6, F1 and H9 derived hPSC-MSNs at DIV 20, 30 and 40 of independent MSN differentiations (n=3 differentiations per cell line). Data is normalised to DIV 30 H9 derived MSNs. Significant results are depicted on bar charts where £ symbolises main effects of line, $ symbolises main effects of Day, # symbolises interactions between Line/Day and * symbolises interactions between Day/Line. The number of symbols indicates p value, where 1 symbol = p<0.05, 2 = p<0.01, and 3 = p<0.001.

