## Supplementary Table 1 for "Defining the Limits of hPSC Derived Models: hPSC-MSNs Fail to Recapitulate Authentic Striatal Identity"

Supplementary Table 1: Key studies of epigenetic memory in iPSCs

| Study | Animal and Tissue of Origin | Did epigenetic memory: | | Summary |
| --- | --- | --- | --- | --- |
|  |  | persist in iPSCs? | effect differentiation? |  |
| Kim *et al.*, 2010 | Mouse.  Bone marrow progenitors (B-iPSC), and dermal fibroblasts (F-iPSC) | Yes  (direct evidence from DNA methylation analysis) | Yes | First systematic study of epigenetic memory in iPSCs. Generated four lines: B-iPSC, F-iPSCs, ESCs, and nuclear transfer stem cells (N-ESC). Found B-iPSCs more readily formed haematopoietic colonies than F-iPSCs, whereas F-iPSCs more readily formed osteogenic colonies. ESC and N-ESCs were as efficient as each other. DNA methylation revealed three main clusters of cells separating B-iPSCs, F-iPSCs, and ESCs and N-ESCs. |
| Polo *et al.*, 2010 | Mouse.  Tail tip fibroblasts, splenic B cells (S-iPSC), bone marrow granulocytes, and skeletal muscle precursors | Yes  (direct evidence from DNA methylation analysis) | Yes | First evidence in truly isogenic lines, as iPSCs were all derived from an iPSC mouse chimera with doxycycline inducible OSKM genes. Though four lines were used, they were run in pairs in separate experiments. iPSCs derived from different lines expressed genes related to their cell type of origin, this was found to correspond to DNA methylation, and a lines propensity to differentiate towards cells of related origin (e.g. S-iPSCs readily differentiated towards macrophage phenotypes). |
| Ghosh *et al.*, 2010 | Human.  Foreskin fibroblast, adipose stem cells, neonatal fibroblast, and keratinocytes | *Yes*  (inferred from indirect evidence) | Not tested | First evidence in human, though differentiation capacity was untested. Transcriptional analysis revealed gene expression was more similar between hiPSCs and their tissue of origin, than all other lines and a hESC line. However, between hESCs and hiPSCs gene expression was consistently more similar than between hiPSCs and their tissue of origin. Of all the hiPSCs, foreskin fibroblasts were most similar to hESCs. No differentiation was conducted. |
| Tian *et al.*, 2011 | Mouse.  Astrocytes (A-iPSC) and fibroblasts (F-iPSC) | *Yes*  (inferred from indirect evidence) | Yes | First evidence in neuroectoderm derived cells. A-iPSCs displayed slower EB formation and reduced proliferation than F-iPSCs and ESCs. However, when differentiating towards a neuronal and dopaminergic fate, A-iPSCs expressed significantly more βIII-tubulin and TH than F-iPSCs. |
| Ohi *et al.*, 2011 | Human.  Hepatocytes, fibroblasts, and melanocytes. | Yes  (direct evidence from DNA methylation analysis) | No  (unpublished data) | Provisional evidence that partially reprogrammed DNA methylation could be required for successful induction to pluripotency rather than random, as knock down of one of these genes reduced reprogramming efficiency. Otherwise, similar findings to earlier studies, hiPSCs retain a small degree of transcription and DNA methylation based memory to their tissue of origin. Differentiation was carried out on the cells, but no differences were observed. |

| Bar-Nur *et al.*, 2011 | Human.  β-pancreatic (β-cell), non-β pancreatic, and fibroblast | Yes  (direct evidence from DNA methylation and histone analysis) | Yes | First evidence that epigenetic memory also influences differentiation in hiPSCs, also suggest that this might be because the differentiation protocols are insufficient in hESCs, and are therefore enhanced by epigenetic memory. Demonstrated β-cell derived hiPSCs maintained more open chromatin structures at critical β-cell genes, and expressed these genes significantly more than all other tested hiPSCs and hESCs both *in vitro* and *in vivo*. |
| --- | --- | --- | --- | --- |
| Xu *et al.*, 2012 | Mouse.  Ventricular myocytes (V-iPSC) and fibroblasts (F-iPSC) | Yes  (direct evidence from DNA methylation analysis) | Yes | Found that during chimera formation, V-iPSCs exhibited a bias to contribute to heart formation, due to their epigenetic memory, and spontaneously formed beating cardiomyocytes 2 days before F-iPSC and ESC controls. Further, they exhibited a bias towards ventricular myocyte differentiation in protocols that typically result in mixed ventricular/atrial myocyte populations. |
| Hargus *et al.*, 2014 | Human.  Fetal neural stem cells (NS-iPSC), cord blood (CB-iPSC), and fibroblast (F-iPSC) | Yes  (direct evidence from DNA methylation analysis) | Yes | First evidence that hiPSCs derived from fetal brain tissue retain an epigenetic memory. NS-iPSCs were enriched for neural genes and retained a methylation signature more similar to their tissue of origin than CB-iPSCs and F-iPSCs. Differences were observed when transplanting *in vivo* to the mouse cortex, indicating improved graft survivability of NS-iPSCs. |
| Phetfong et al., 2016 | Human.  Umbilical cord vein endothelial cells (UE-iPSC), endothelial progenitors (E-iPSC), fibroblasts (F-iPSC) | Yes  (direct evidence from DNA methylation analysis) | Yes | Found that UE-iPSCs and E-iPSCs both had improved potentiality towards endothelial cell differentiation compared to F-iPSCs. Only UE-iPSCs exhibited efficient differentiation towards a haematopoietic progenitor phenotype. Additionally, both UE- and E-iPSCs showed increased efficiency of initial iPSC reprogramming. DNA methylation differences were shown at key endothelial genetic regions. |
| Roost *et al.,* 2017 | Human.  Fetal brain, skin, kidney, muscle, lung, pancreas. | Yes  (direct evidence from DNA methylation analysis) | Yes | Comprehensive study exploring methylome of 21 human fetal organs across fetal development, including brain tissue. Generated isogenic hiPSCs from 6 organs, found hiPSCs derived from fetal brain were more capable of differentiating into neural phenotypes compared to non-neural derived hiPSC controls. |
| Khan *et al.,* 2023 | Human.  Healthy and Osteoarthritic chondrocytes. | Yes  (direct evidence from DNA methylation analysis) | Yes | Compared hiPSCs derived from healthy and unhealthy (osteoarthritic) chondrocytes, finding those derived from healthy are more capable of re-differentiating into chondrocytes, suggesting disease state may also influence resulting hiPSC fate commitment. |
