## Supplementary Table 2 for "Defining the Limits of hPSC Derived Models: hPSC-MSNs Fail to Recapitulate Authentic Striatal Identity"

Supplementary Table 2 – DNA methylation sample information

| **Sample name** | **Genetic identity** | **Cell/tissue phenotype** | **% of failed probes** |
| --- | --- | --- | --- |
| Primary LGE 2415 | SWIFT 2415 | Fetal LGE | 0.1% |
| Primary LGE 2451 | SWIFT 2451 | Fetal LGE | 0.01% |
| Primary LGE 2285 | **SWIFT 2285** | Fetal LGE | 0.06% |
| Primary FIB 2285 | **SWIFT 2285** | Fetal fibroblast | 0.05% |
| hiPSC L1 Pluri | **SWIFT 2285** | hiPSC | 0.06% |
| hiPSC L2 Pluri | **SWIFT 2285** | hiPSC | 0.17% |
| hiPSC L11 Pluri | **SWIFT 2285** | hiPSC | 0.06% |
| hiPSC F1 Pluri | **SWIFT 2285** | hiPSC | 0.08% |
| hESC H9 Pluri | ESC H9 | hESC | 0.05% |
| hiPSC L1 MSN i | **SWIFT 2285** | hiPSC-MSN | 0.06% |
| hiPSC L1 MSN ii | **SWIFT 2285** | hiPSC-MSN | 0.05% |
| hiPSC L1 MSN iii | **SWIFT 2285** | hiPSC-MSN | 0.11% |
| hiPSC L2 MSN i | **SWIFT 2285** | hiPSC-MSN | 0.72% |
| hiPSC L2 MSN ii | **SWIFT 2285** | hiPSC-MSN | 0.06% |
| hiPSC L2 MSN iii | **SWIFT 2285** | hiPSC-MSN | 0.50% |
| hiPSC L11 MSN i | **SWIFT 2285** | hiPSC-MSN | 0.36% |
| hiPSC L11 MSN ii | **SWIFT 2285** | hiPSC-MSN | 0.05% |
| hiPSC L11 MSN iii | **SWIFT 2285** | hiPSC-MSN | 0.06% |
| hiPSC F1 MSN i | **SWIFT 2285** | hiPSC-MSN | 0.06% |
| hiPSC F1 MSN ii | **SWIFT 2285** | hiPSC-MSN | 0.31% |
| hiPSC F1 MSN iii | **SWIFT 2285** | hiPSC-MSN | 0.06% |
| hESC H9 MSN i | ESC H9 | hESC-MSN | 0.05% |
| hESC H9 MSN ii | ESC H9 | hESC-MSN | 0.05% |
